# Weight Sensitivity of Temporal SNR Metrics in multi-echo fMRI

**DOI:** 10.1101/2020.04.20.049460

**Authors:** Thomas T. Liu, Bochao Li, Conan Chen, Brice Fernandez, Baolian Yang, Suchandrima Banerjee

## Abstract

**Purpose:** In multi-echo fMRI (ME-fMRI), various weighting schemes have been proposed for the combination of the data across echoes. Here we introduce a framework that facilitates a deeper understanding of the weight dependence of temporal SNR measures in ME-fMRI.

**Theory and Methods:** We examine two metrics that have been used to characterize ME-fMRI performance: temporal SNR (tSNR) and multi-echo temporal (metSNR). Both metrics can be described using the generalized Rayleigh quotient (GRQ) and are predicted to be relatively insensitive to the weights when there is a high degree of similarity between a metric-specific matrix in the GRQ numerator and a metric-independent covariance matrix in the GRQ denominator. The application of the GRQ framework to experimental data is demonstrated using a resting-state fMRI dataset acquired with a multi-echo multi-band EPI sequence.

**Results:** In the example dataset, similarities between the covariance matrix and the metSNR and tSNR numerator matrices are highest in grey matter (GM) and cerebrospinal fluid (CSF) voxels, respectively. For representative GM and CSF voxels that exhibit high matrix similarity values, the metSNR and tSNR values, respectively, are both within 4% of their optimal values across a range of weighting schemes. However, there is a fundamental tradeoff, with a high degree of weight sensitivity in the tSNR and metSNR metrics for the representative GM and CSF voxels, respectively. Geometric insight into the observed weight dependencies is provided through a graphical interpretation of the GRQ.

**Conclusion:** A GRQ framework can provide insight into the factors that determine the weight sensitivity of tSNR and metSNR measures in ME-fMRI.

## Introduction

A key step in multi-echo fMRI (ME-fMRI) is the weighted combination of the data from multiple echoes. Various weighting schemes have been proposed [1, 2, 3], and the contrast-to-noise (CNR) values achieved by the different methods have been found to be relatively similar. In this technical note, we consider two metrics that have been used to characterize the CNR of multi-echo fMRI: temporal SNR (tSNR) and multiecho temporal SNR (metSNR) [2, 3, 4]. We first demonstrate the relation between these metrics and the generalized Rayleigh quotient (GRQ). We then show how the framework of the GRQ can be used to provide insights into the relative sensitivity of the metrics to the choice of weights. A preliminary version of this work was presented in [5].

## Theory

For *N*_*E*_ echoes, we define **w** as the *N*_*E*_ × 1 vector of weights and **S** as a *N*_*E*_ × *N*_*T*_ matrix where the *i*th row is the time series data from the *i*th echo. With this notation, **w**^*T*^**S** represents the weighted combination of data and

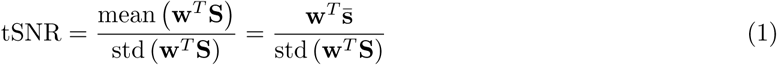

where 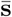 is the *N*_*E*_ × 1 vector consisting of the temporal means (i.e. row means of **S**). To maximize tSNR, we can consider maximizing its square

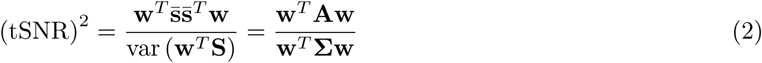

where 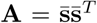 and **Σ** denotes the covariance matrix of **S**. The expression for (tSNR)^2^ has the form of a GRQ and attains a maximum value of 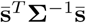 when the weight vector has the form of an optimal matched filter 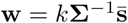, where *k* is an arbitrary scalar [6, 7, 8]. From the form of the GRQ, it is straightforward to show that 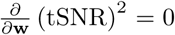 when **A** is a scalar multiple of **S**. Thus, we expect that tSNR will be relatively insensitive to the form of **w** when **A** ∼ **Σ**.

For a single echo acquisition, tSNR is proportional to CNR. For multi-echo acquisitions, a multi-echo variant of tSNR that is proportional to CNR [2, 3] is defined as:

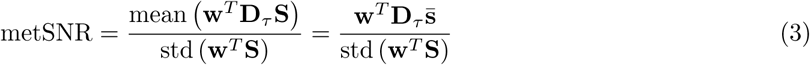

where 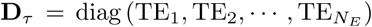 is the diagonal matrix comprised of the echo times. The corresponding GRQ is

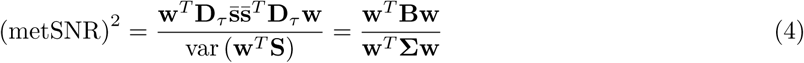

where 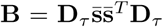. The GRQ attains its maximal value 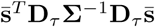 when 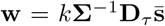. Further-more, the GRQ and hence metSNR are both fairly insensitive to the choice of weights when **B** ∼ Σ. Since both tSNR and metSNR are invariant with respect to the value of *k*, we will drop this scalar term for the remainder of the paper. Background material on the relation between tSNR, metSNR, and CNR is provided in the Appendix.

Under certain conditions, previously proposed weighting schemes can achieve optimal tSNR or metSNR. For example, with the assumption that the random noise components have equal variance and are uncorrelated across the echoes (i.e. Σ = *σ*^2^**I**), the optimal weight vector for metSNR has the form 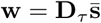, which is the BOLD Sensitivity weight solution examined in [3]. This is equivalent to the matched filter solution proposed in [1] when the signal means exhibit 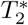 weighting (i.e. 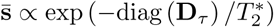), as would be the case for most ME-fMRI acquisitions. For tSNR maximization, the assumption Σ = *σ*^2^**I** leads to an optimal weight vector of the form 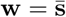, which is analogous to the exponential weighting considered by [1].

If the noise is uncorrelated with unequal variances across echoes (i.e. 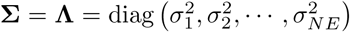), then the optimal metSNR weight vector has the form 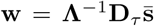. This solution is similar to but not equivalent to the temporal BOLD sensitivity weighting 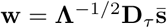 proposed in [2] and further examined in [3]. For tSNR maximization, the optimal weight vector when **Σ** = **Λ** is 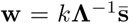, which is similar to but not equivalent to the tSNR weighting 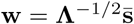 studied in [3].

The various options described above for the weight vector are summarized in Table 1. The first four options consist of the tSNR optimal solution followed by three variations listed in order of decreasing similarity to the optimal solution. The last five options consist of the metSNR optimal solution followed by three variations listed in order of decreasing similarity to the optimal solution. In addition, we also consider a flat weighting of the echoes that has been examined in prior studies [1, 3, 9].

**Table 1:**
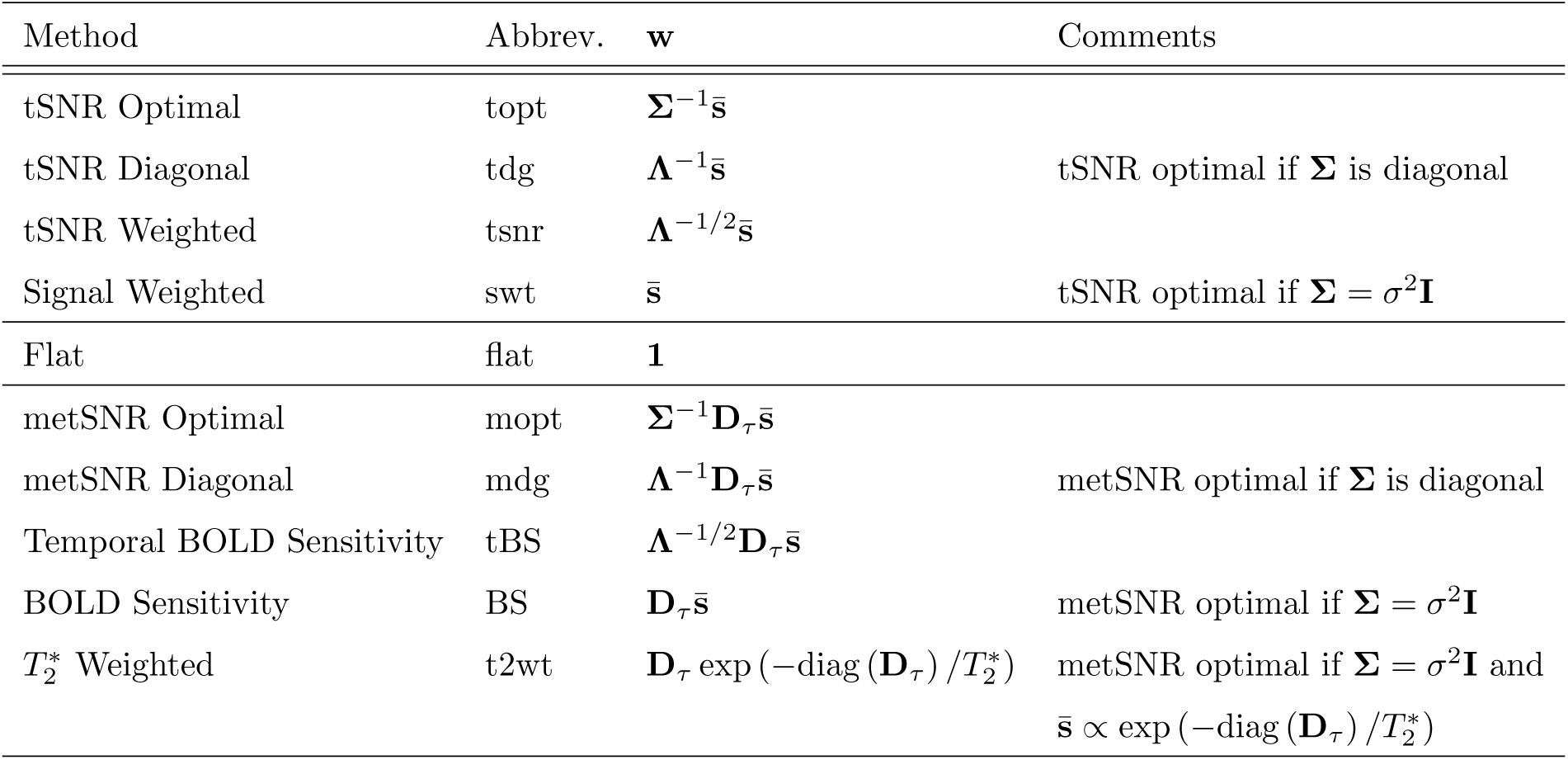
Description of weight vectors.

| Method | Abbrev. | $\mathbf{w}$ | Comments |
| --- | --- | --- | --- |
| tSNR Optimal | topt | $\mathbf{\Sigma}^{-1}\bar{\mathbf{s}}$ | |
| tSNR Diagonal | tdg | $\mathbf{\Lambda}^{-1}\bar{\mathbf{s}}$ | tSNR optimal if $\mathbf{\Sigma}$ is diagonal |
| tSNR Weighted | tsnr | $\mathbf{\Lambda}^{-1/2}\bar{\mathbf{s}}$ | |
| Signal Weighted | swt | $\bar{\mathbf{s}}$ | tSNR optimal if $\mathbf{\Sigma} = \sigma^2\mathbf{I}$ |
| Flat | flat | $\mathbf{1}$ | |
| metSNR Optimal | mopt | $\mathbf{\Sigma}^{-1}\mathbf{D}_\tau\bar{\mathbf{s}}$ | |
| metSNR Diagonal | mdg | $\mathbf{\Lambda}^{-1}\mathbf{D}_\tau\bar{\mathbf{s}}$ | metSNR optimal if $\mathbf{\Sigma}$ is diagonal |
| Temporal BOLD Sensitivity | tBS | $\mathbf{\Lambda}^{-1/2}\mathbf{D}_\tau\bar{\mathbf{s}}$ | |
| BOLD Sensitivity | BS | $\mathbf{D}_\tau\bar{\mathbf{s}}$ | metSNR optimal if $\mathbf{\Sigma} = \sigma^2\mathbf{I}$ |
| $T_2^*$ Weighted | t2wt | $\mathbf{D}_\tau \exp(-\text{diag}(\mathbf{D}_\tau)/T_2^*)$ | metSNR optimal if $\mathbf{\Sigma} = \sigma^2\mathbf{I}$ and<br>$\bar{\mathbf{s}} \propto \exp(-\text{diag}(\mathbf{D}_\tau)/T_2^*)$ |

## Methods

To demonstrate the application of the theory to experimental data, we acquired resting-state fMRI data (eyes-closed) under Institutional Review Board approval. One healthy male was scanned on a Discovery MR750 3T system (GE Healthcare) with a 32-channel receive head coil (Nova Medical) and a multi-echo multi-band EPI sequence with the following parameters: TR = 1.3 s; *θ* = 52°; FOV = 192 mm; matrix size = 64 × 64; 48 3 mm thick slices, multi-band factor = 3; TE=[12.2 30.1 48.0] ms, 277 reps; scan time = 6 min. Data were preprocessed using the meica.py script (v2.5b11) with 2nd order polynomial detrending, default despiking, and no smoothing [4]. For each voxel we computed values of tSNR and metSNR, using Equations 1 and 3 and the weight vectors (with sample means and covariances) listed in Table 1.

## Results

### tSNR and metSNR maps

Figure 1 shows tSNR and metSNR maps obtained using the different weight vectors, labeled using the abbreviations and ordering from Table 1. For each voxel, the tSNR and metSNR values are normalized by the maximum values of 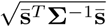 and 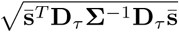, respectively, that are obtained when using the optimal weight vector for each metric.

**Figure 1:**
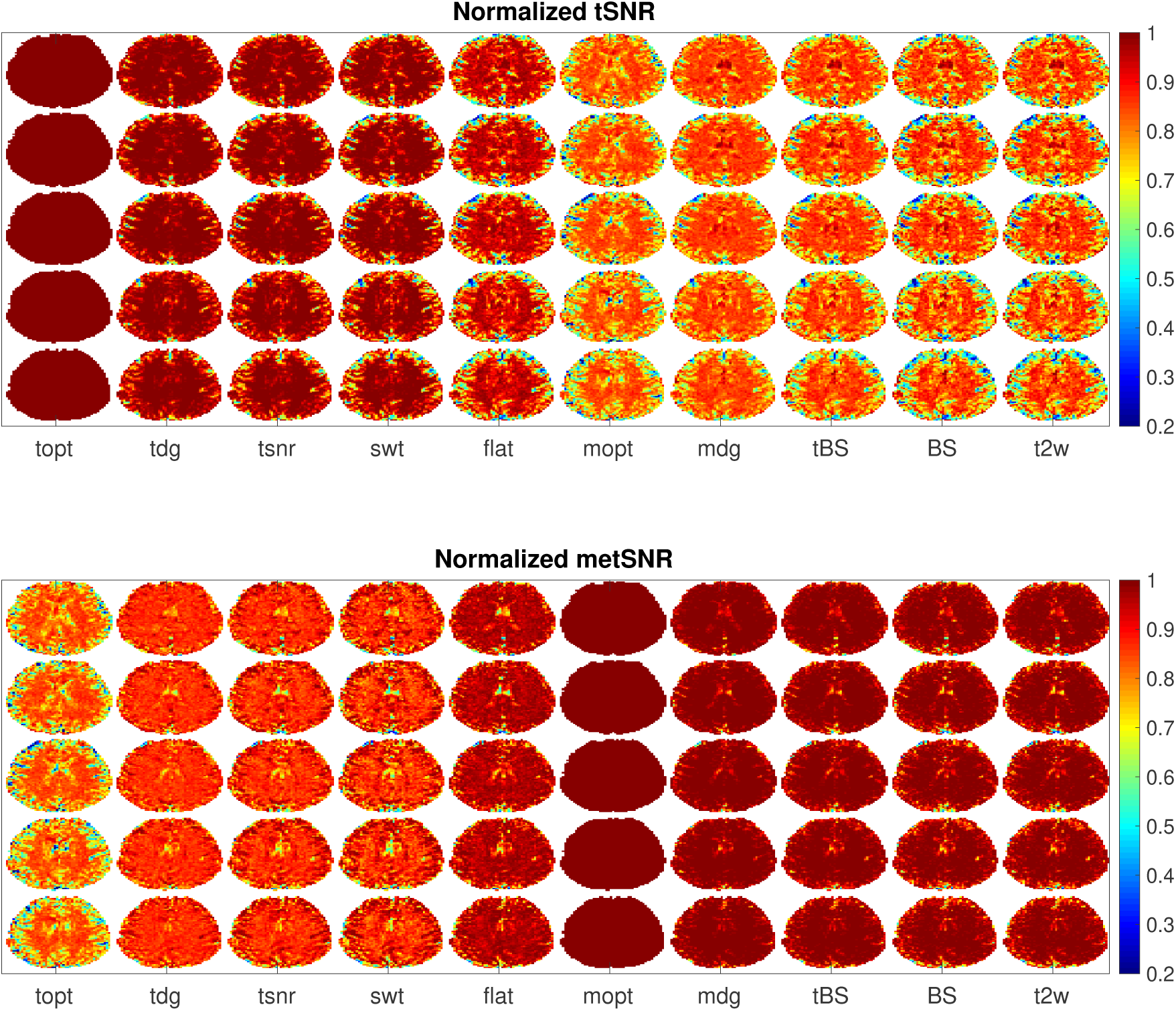
Maps of tSNR and metSNR for 5 slices and 10 different weight vectors. Columns are labeled using the abbreviations listed in Table 1. Maps are normalized by the values obtained using their respective optimal weights, such that the maximum value is 1.0. Thus, the tSNR and metSNR maps obtained using the tSNR and metSNR optimal weights (topt and mopt), respectively, are both uniformly equal to 1.0, while maps using other weight vectors have values less than 1.0.

The maps can be roughly split into (1) a group with higher tSNR and lower metSNR values obtained using weight vectors (topt, tdg, tsnr, and swt) that are variations of the tSNR optimal weight vector (topt) and (2) a group with lower overall tSNR and higher metSNR values obtained using weight vectors (mopt, mdg, tBS, BS, t2w) that are variations of the metSNR optimal weight vector (mopt). The tSNR and metSNR maps obtained with the flat weight vector lie between these two groups.

Note that the normalized tSNR maps obtained with the metSNR optimal weights (column mopt in the upper part of the Figure) are identical to the normalized metSNR maps obtained with the tSNR optimal weights (column topt in the lower part of the Figure). Indeed, it is straightforward to show that

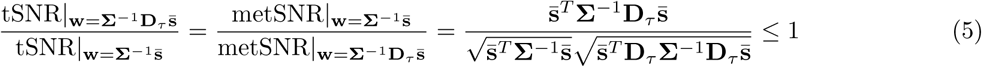

where the inequality follows from the Cauchy-Schwarz inequality.

### Similarity between matrices

As noted in the Theory section, tSNR and metSNR are expected to be relatively insensitive to the choice of weights when the respective numerator matrices (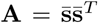 and 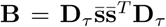) are similar in form to the covariance matrix Σ. As a measure of similarity between matrices, we computed the cosine similarity for each voxel using the six unique terms of each matrix. The first row of Figure 2 shows that the cosine similarity between **A** and **Σ** is relatively high in the cerebral spinal fluid (CSF) of the lateral ventricles and superficial cortical gyri. In contrast, the cosine similarity between **B** and **Σ** (second row) is relatively high in superficial cortical and deep grey matter (GM) regions. The third row shows that the cosine similarity between **Σ** and the identity matrix (i.e. **Σ** ∼ **I**) is relatively high in white matter voxels. Representative GM, CSF, and WM voxels are indicated in the maps.

**Figure 2:**
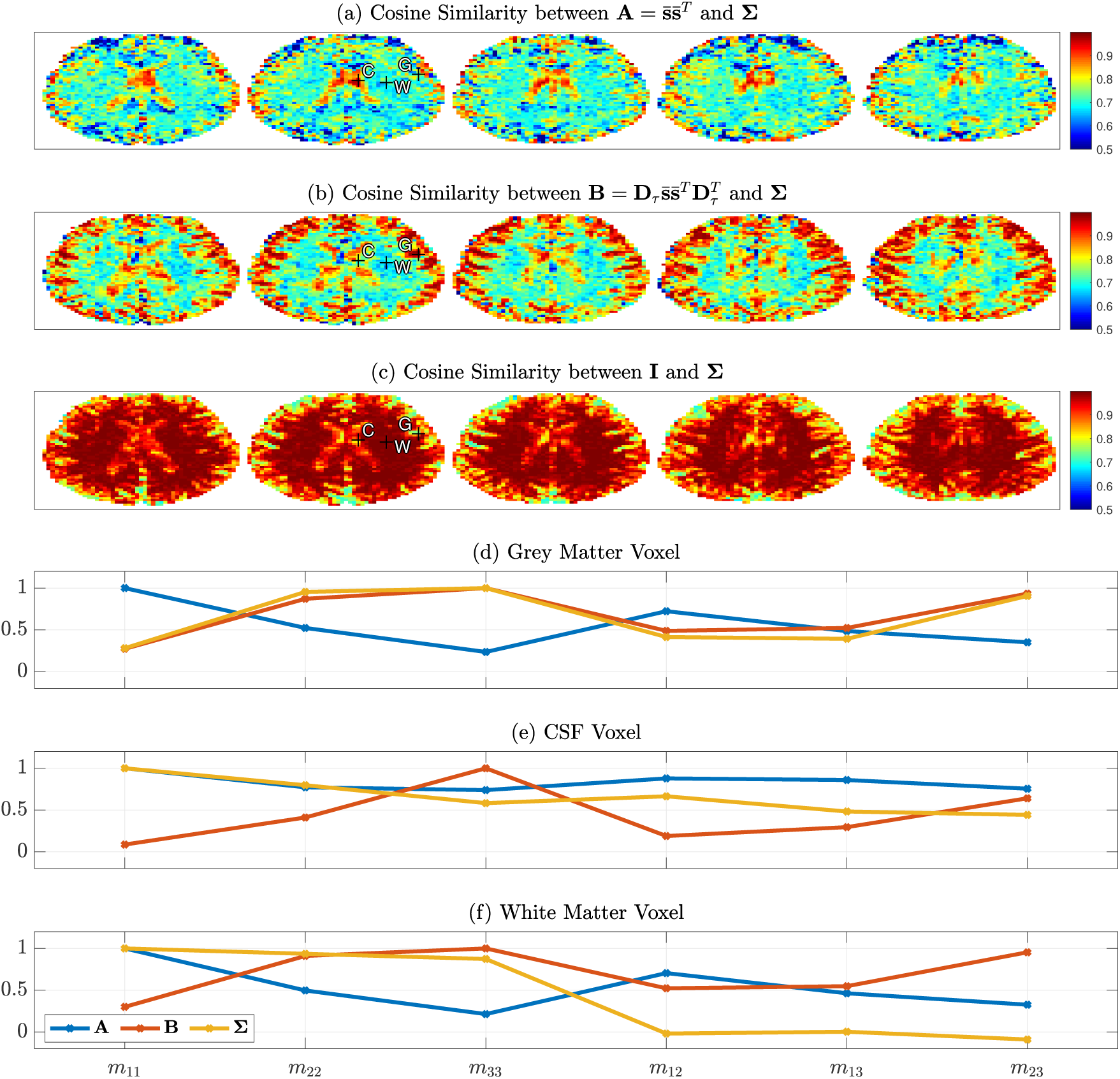
(a-c) Maps of cosine similarity between the covariance matrix **Σ** and tSNR numerator matrix **A**, metSNR numerator matrix **B**, and the identity matrix **I**. Representative voxels in grey matter, cerebrospinal fluid (CSF), and white matter regions are marked with plus symbols and the letters G, C, and W, respectively. (d-f) Matrix elements *m*_*ij*_ of the matrices **A**, **B**, and **Σ** plotted for each of the representative voxels, where the indices *i* and *j* refer to the echo number, *m*_11_, *m*_22_, and *m*_33_ are the diagonal terms, and *m*_12_, *m*_13_, and *m*_23_ are the off-diagonal terms. For each matrix, the values are normalized by the maximum value.

The lower half of Figure 2 plots the elements of the matrices **A**, **B**, and **Σ** for each of the representative voxels, with matrix values normalized by their respective maximum value. For the GM voxel, the diagonal elements (*m*_11_ through *m*_33_) of both the **B** and **Σ** matrices show an increase with echo time and the offdiagonal term is highest for the interaction (*m*_23_) between the second and third echoes. In contrast, both the diagonal elements and interaction terms of **A** decrease with echo time. As a result, the cosine similarity between **B** and **Σ** has a value (0.995) close to the maximum of 1.0 while the similarity between **A** and **Σ** has a value of 0.693. The fact that **Σ** ∼ **B** indicates that the observed noise in each echo is roughly “BOLD-like” [9]. Due to the high degree of similarity between **B** and Σ, the metSNR values (Figure 3a) are relatively insensitive to the choice of weight vector (ranging from 0.96 to 1.0), with the exception of the noticeably lower value of 0.54 for the tSNR optimal weight. In contrast, the tSNR values exhibit a greater degree of sensitivity with values ranging from 0.34 to 1.0.

**Figure 3:**
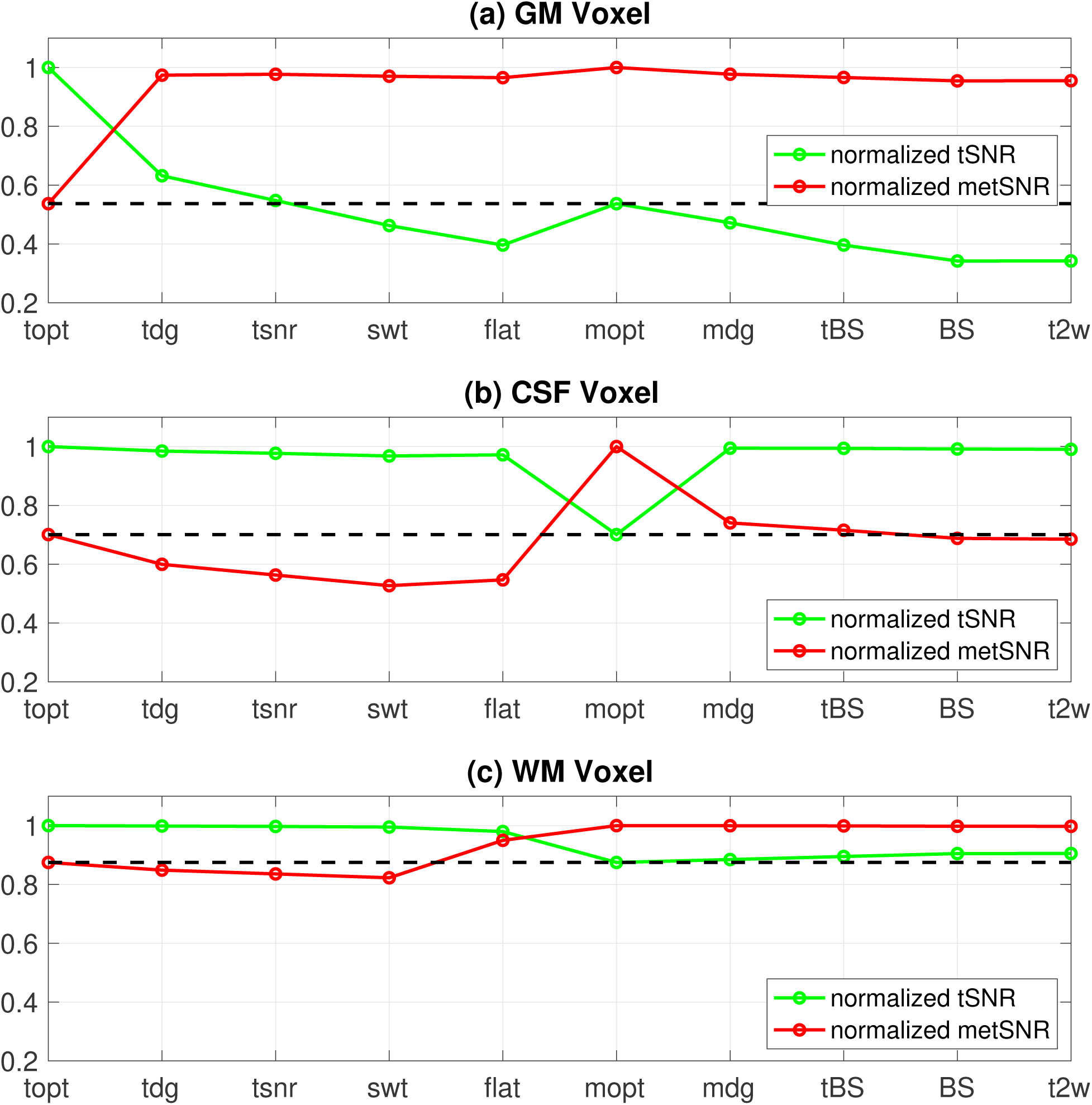
Normalized tSNR and metSNR values plotted versus weight vector for the representative voxels indicated in Figure 2. Weight vectors are labeled with the abbreviations listed in Table 1. The black dotted lines are provided to point out that the normalized tSNR and metSNR values obtained when using metSNR and tSNR optimal weights, respectively, have the same value.

For the CSF voxel the diagonal elements and interaction terms of both **A** and Σ decrease with echo time, whereas these terms increase with echo time in **B**, resulting in corresponding similarity values of 0.975 and 0.700. The high similarity between **A** and **Σ** indicates that the noise in each echo is roughly “signal-like” [9]. Due to the high degree of similarity between **A** and **Σ**, the tSNR values shown in Figure 3b are relatively insensitive to the choice of weight vector (ranging from 0.97 to 1.0), with the exception of the noticeably lower value of 0.70 for the metSNR optimal weight. In contrast, the metSNR values exhibit a greater degree of sensitivity with values ranging from 0.53 to 1.0.

The covariance matrix **Σ** for the WM voxel has diagonal elements that decrease slightly with echo time and off-diagonal terms that are close to zero, exhibiting a high degree of simlarity (0.997) with the identity matrix. In contrast, the mismatch between **Σ** and both **A** and **B** is reflected in cosine similarity values of 0.682 and 0.644, respectively. Despite these low similarity values, the tSNR and metSNR values shown in Figure 3c for this voxel show a moderate to high degree of insensitivity to the choice of weight vector. The normalized tSNR values for the tSNR optimal weight variants (topt, tdiag, tsnr, swt) are within 0.005 of the maximum value of 1.000, reflecting the fact that all of the variants approach the optimal solution 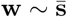 when **Σ** ∼ **I**. Similarly, the metSNR values for the metSNR optimal weight variants (mopt, mdg, tBS, BS, t2wt) are within 0.003 of the maximum value of 1.000, reflecting the fact that all of these variants approach the optimal solution 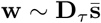.

### Geometric Intepretation

To better understand the weight sensitivity, it is useful to consider a geometric representation. In Figure 4 the first and last rows show the absolute values of the normalized tSNR and metSNR values, respectively, plotted on the surface of a sphere, where each point of the sphere corresponds to a weight vector with unit norm. The second and fourth rows show the absolute values of the tSNR 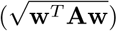 and metSNR 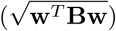 numerator terms, respectively, while the third row show the common denominator term 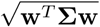. As an alternate way of viewing the quantities, Figure 5 shows polar plots in which the radius is proportional to the normalized quantity of interest.

**Figure 4:**
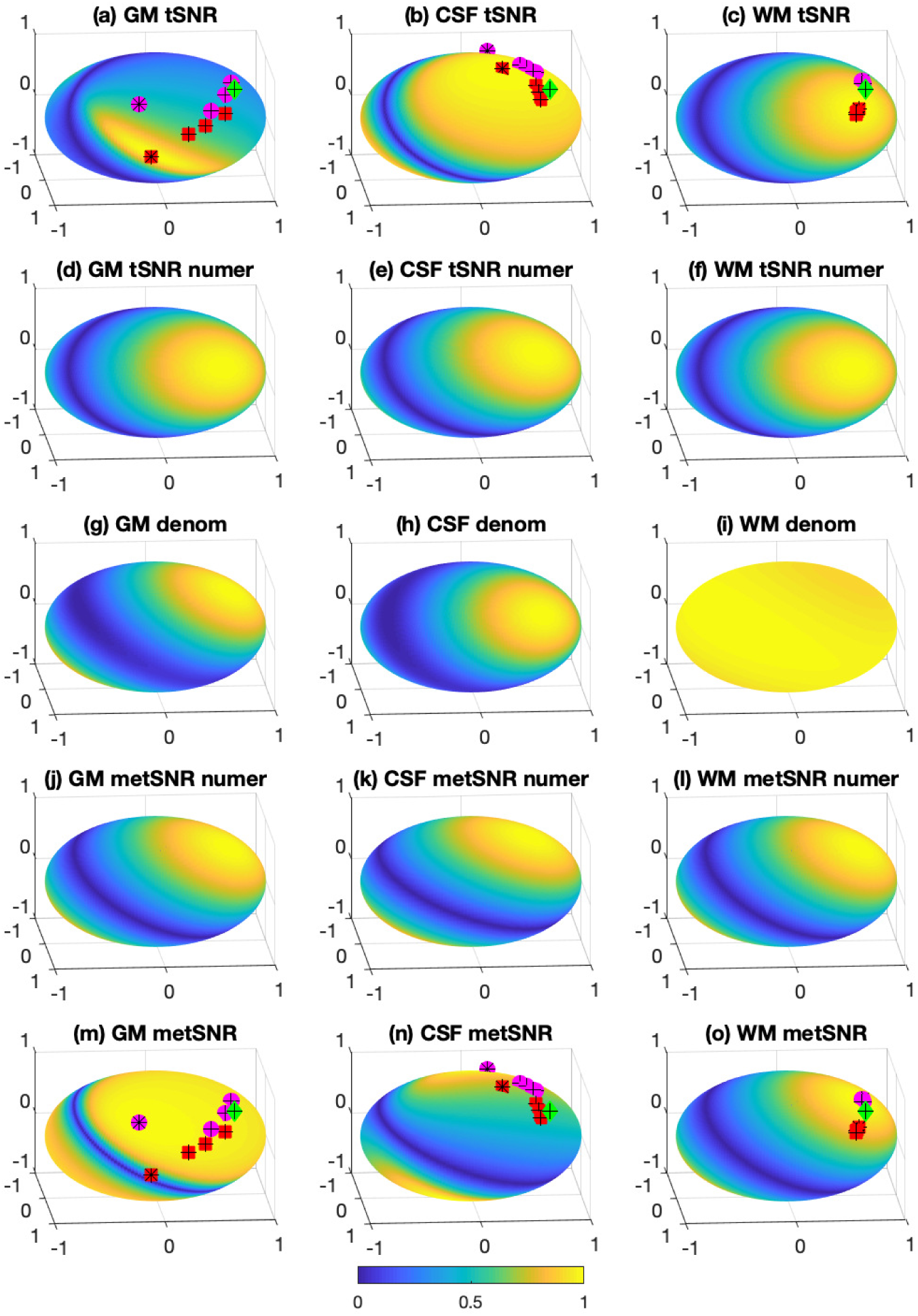
Spherical surface plots showing the weight dependence of (a-c) tSNR, (d-f) tSNR numerator, (g-i) denominator, (j-l) metSNR numerator, and (m-o) metSNR for each of the representative voxels (GM, CSF, and WM organized by columns from left to right) indicated in Figure 2. All quantities are shown as absolute values and normalized by their maximum value. The red squares indicate tSNR weight variants (topt, tdg, tsnr, swt), the magenta circles indicates metSNR weight variants (mopt, mdg, tBS, BS, t2wt), and the green diamonds indicate the flat weight vectors. The optimal tSNR and metSNR weights are indicated with black asterisks while the other variants have black crosses.

**Figure 5:**
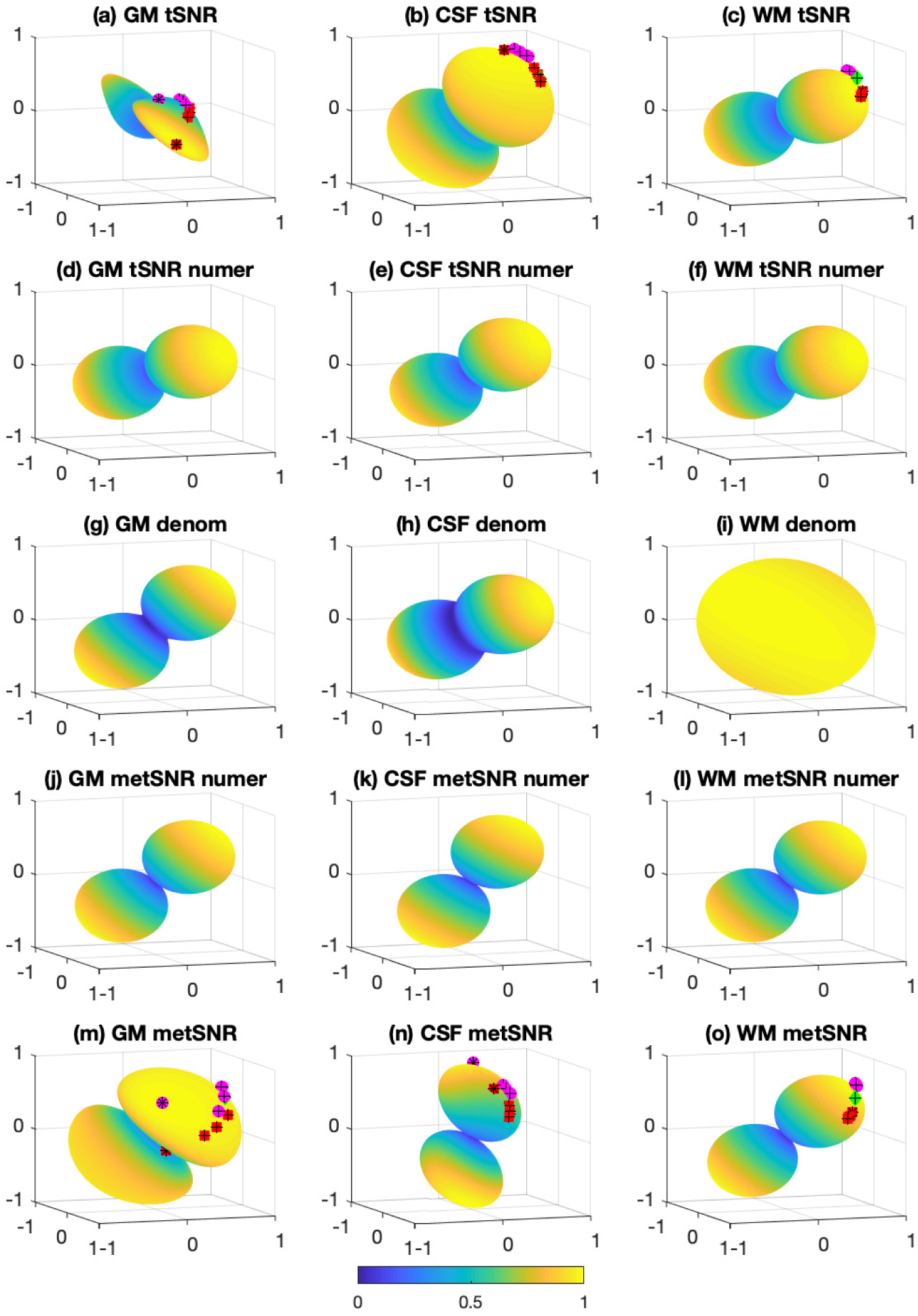
Polar plots showing the weight dependence of (a-c) tSNR, (d-f) tSNR numerator, (g-i) denominator, (j-l) metSNR numerator, and (m-o) metSNR for each of the representative voxels (GM, CSF, and WM organized by columns from left to right) indicated in Figure 2. All quantities are shown as absolute values and normalized by their maximum value. The red squares indicate tSNR weight variants (topt, tdg, tsnr, swt), the magenta circles indicates metSNR weight variants (mopt, mdg, tBS, BS, t2wt), and the green diamonds indicate the flat weight vectors. The optimal tSNR and metSNR weights are indicated with black asterisks while the other variants have black crosses. The optimal metSNR weight variant for the CSF voxel is obscured from view.

All surface plots are normalized to have a maximum value of 1.0 for display purposes. With this normalization, the numerator surface plots represent the functions 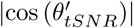 and 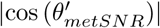 where 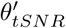 and 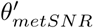 are the polar angles referenced to the vectors 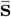 and 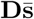 for the tSNR and metSNR numerators, respectively. Since **D**_*τ*_ is independent of voxel, variations across voxels in the numerator terms are driven solely by variations in the mean signal vector 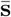, which in turn reflects the 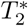 decay in each voxel. The similarity in numerator surface plots thus reflects the first order similarity of the decay curves across tissue types.

The tSNR and metSNR surface plots are obtained by taking the ratio of the respective numerator plot and the common denominator plot. Since the numerator plots are fairly similar across tissue types, the voxel-wise differences in the tSNR and metSNR surface plots are driven primarily by differences in the denominator term, which reflect differences in **Σ**.

For the GM voxel, the metSNR numerator (panel j) and denominator (panel g) plots exhibit similar dependencies on the weights, reflecting the fact that **Σ** ∼ **B**. Further insight can be gained by noting that **Σ** has a dominant eigenvector **v**_1_, so that its corresponding surface plot can be approximated as 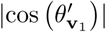 where 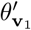 is the polar angle referenced to **v**_1_. Because the angle between **v**_1_ and 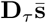 is relatively small (∆*θ* = 3.5°), the numerator and denominator plots exhibit similar cosine dependencies on the weights. As a result, the ratio of the two plots results in a metSNR surface plot (panel m) that is relatively insensitive to **w**. In contrast, the principal directions of the tSNR numerator (panel d) and denominator (panel g) plots are not well aligned (∆*θ* = 32.6°), so that the ratio of these two plots results in a tSNR surface plot (panel a) that is sensitive to **w**.

For the CSF voxel, the tSNR numerator (panel e) and denominator (panel h) plots exhibit similar dependencies on the weights, reflecting the fact that **Σ** ∼ **A** with a dominant eigenvector that is nearly aligned (∆*θ* = 6.3°) with the vector 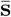. As a result, the tSNR surface plot (panel b) is relatively insensitive to **w**. On the other hand, the principal directions of the metSNR numerator (panel k) and denominator (panel h) are not well aligned (∆*θ* = 33.4°), so that the ratio of these two terms results in a metSNR surface plot (panel n) that is sensitive to **w**.

For the WM voxel, the denominator plot (panel i) is fairly uniform, reflecting the fact that **Σ** ∼ **I** with a fairly uniform spread of eigenvalues (normalized range of 1.0 to 1.3). As a result, the tSNR (panel c) and metSNR (panel o) surface plots primarily reflect the weight dependencies of their respective numerator terms (panels f and l). In contrast to the spread in weight vector positions observed for the GM and CSF voxels, the WM weight locations are fairly tightly clustered since, as noted above, the tSNR and metSNR weight variants approach the optimal solutions 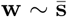 and 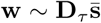, respectively. The tSNR and metSNR surface plots reach their maxima near these weight vectors, with red squares clustered around 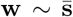 in panel c and magenta circles clustered around 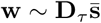 in panel o. Furthermore, because of the approximate cosine dependence of the surface plots, the tSNR and metSNR values obtained when using metSNR and tSNR weight variants (magenta circles in panel c and red squares in panel o) can be approximated with the cosine of the angle between 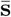 and 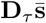. For the WM voxel, 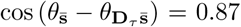 which is equal to the mean of the relevant values (metSNR values for topt, tdg, tnsr, and swt and tSNR values for mopt, mdg, tBS, BS, and t2wt; range of 0.83 to 0.91) shown in Figure 3(c).

## Discussion and Conclusions

We have shown that tSNR and metSNR can be described with the GRQ, which is maximized with an optimal matched filter that uses an estimate of the covariance matrix **Σ**. For metSNR, the optimal solution is a generalization of previously proposed matched filter variants [1, 2], which have assumed restrictions on the form of **Σ**. Because the optimal solutions can attain the maximum values of tSNR and metSNR they are useful for assessing the relative performance of other weighting schemes.

The weight sensitivity of tSNR and metSNR depends on the structure of the covariance matrix **Σ** and its similarity with either the respective numerator matrices **A** and **B** or with the identity matrix **I**. As **A** and **B** have different forms, there is a fundamental tradeoff wherein a high similarity between **B** and **Σ** results in metSNR metrics with low weight sensitivity and tSNR metrics with sensitivity. On the other hand, a high similarity between **A** and **Σ** results in tSNR metrics that are relatively insensitive and metSNR metrics that are highly sensitive. Representative voxels in GM and CSF regions were used to demonstrate the link between the similarity in matrices and the corresponding weight sensitivities and tradeoffs. For WM voxels, the high degree of similarity between **Σ** and the identity matrix leads to similar within-group performance for both tSNR and metSNR weight variants. A graphical interpretation of the GRQ was introduced to provide geometric insights into the weight dependencies and the inherent tradeoffs.

Although tSNR has been used to assess ME-fMRI performance [4, 10, 11], it is strictly proportional to CNR only when data are acquired at a single echo time [2]. The weighting schemes (t2wt and flat) used in these prior studies are likely to achieve poor tSNR performance in GM voxels as compared to the optimal weight choice, due to the expected mismatch between **A** and **Σ**. While the weight insensitivity of tSNR in CSF is interesting, its practical relevance for most fMRI studies is limited. These factors suggest that researchers may want to minimize the use of tSNR for the characterization of ME-fMRI performance in GM regions.

On the other hand, metSNR is designed to be proportional to CNR for ME-fMRI data [2] and may therefore be more relevant for assessing ME-fMRI performance. In examining metSNR for three weighting schemes, Poser et al [2] found that the mean CNRs of flat and t2wt weighting schemes were about 93% and 96%, respectively, of the mean CNR obtained with tBS, albeit with considerable variance in the metrics. Using a different protocol, Kettinger et al [3] found that the average CNRs of tsnr, flat, and tBS were 86%, 91%, and 94%, respectively, of the CNR with BS weights but concluded that all the weighting schemes provided similar group-level statistical performance. Some of the observed weight sensitivity may be due to partial voluming effects in the regions of interests used (e.g. a relatively lax threshold of 0.35 was used to define the GM mask in [3]), resulting in metSNR values that reflect a mix of the weight sensitivities of different tissue types. Taken as a whole, the prior findings and those of the present work suggest that the choice of weights may have a rather limited effect on metSNR, especially for GM voxels in which the physiological fluctuations are expected to be “BOLD-like” [4, 9].

In conclusion, we have used the GRQ framework to show how ME-fMRI weight sensitivity depends on the structure of **Σ** and its similarity to the matrices **A**, **B**, and **I**. In addition to tissue type, the form of **Σ** and the associated matrix similarities will depend on a number of experimental factors, such as spatial resolution, echo spacing, field strength, and physiology. Future work to more fully assess the matrix similarities and associated weight sensitivities across a range of factors would be of interest.

## Appendix Background Notes

In this section we review the relations between CNR and the tSNR and metSNR metrics that were previously defined and examined in [12, 13, 2]. While this background material is not critical for the conclusions of the paper, it may be helpful for some readers.

For a single echo acquisition, the measure for CNR is

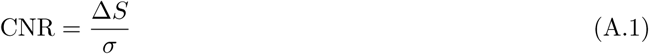

where ∆*S* is the signal change and *σ* is the standard deviation. For fMRI, we typically think in terms of percent BOLD change %∆*S* = 100 × ∆*S/S*(0), where *S*(0) denotes the baseline signal. In practice, the temporal mean of the signal 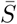 serves as a useful proxy for the baseline signal, so that 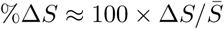. Using this expression and dropping the scaling term 100, we obtain

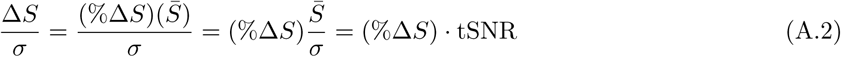

where 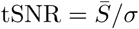. Equation A.2 shows why tSNR is a useful measure. For any given expected %∆*S* (which depends on experimental design, physiology, etc), the CNR will be maximized when tSNR is maximized.

For multiple echo acquisitions, the percent BOLD signal change for the *k*th echo is 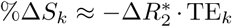[2, 4]. Thus for a given value of 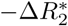, we have %∆*S*_*k*_ ∝ TE_*k*_ and 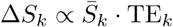.

For the weighted sum of signals 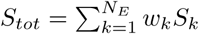, the desired expression for CNR is

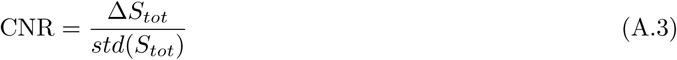

For the numerator of the CNR expression we have

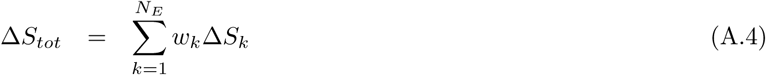

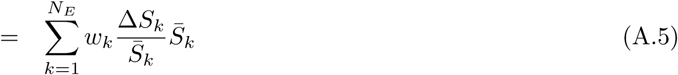

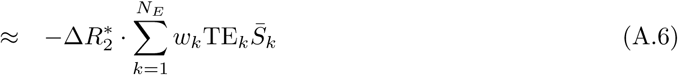

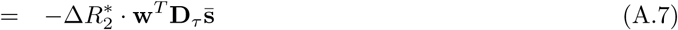

where 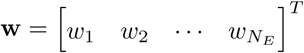, 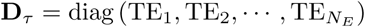, and 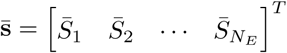. The denominator of the CNR expression is the standard deviation of the weighted signal *std*(**w**^*T*^**S**) where **S** is the matrix with the *k*th row containing the signal from the *k*th echo. Putting everything together, we obtain

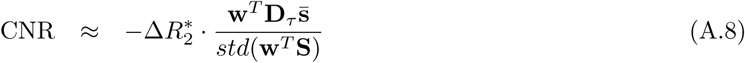

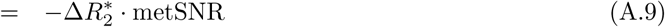

where

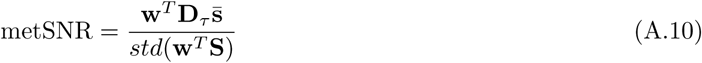

denotes multi-echo tSNR.

The ratio of the CNRs for the single echo (SE) and multi-echo (ME) acquisitions is given by

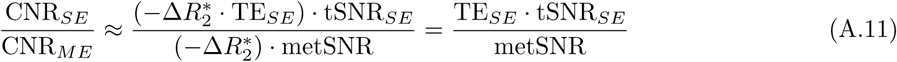

As noted in the main text, metSNR is independent of the weights when **Σ** = *a***B** where *a* is an arbitrary scalar. With this condition 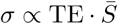, so that TE_*SE*_ ⋅ tSNR_*SE*_ is a constant. Furthermore, it is straightforward to show that TE_*SE*_ · tSNR_*SE*_ = metSNR, so that the CNRs of the single echo and multi-echo acquisitions are equal when **Σ** = *a***B**.

The expression for tSNR in a multi-echo acquisition is

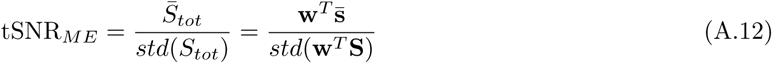

Note that this expression is not in general proportional to the CNR of the multi-echo acquisition. As a sanity check, if all the echo times in the multi-echo acquisition are equal to TE_*SE*_ then TE_*SE*_ ⋅ tSNR_*ME*_ = metSNR and tSNR_*ME*_ = tSNR_*SE*_.

## References

[1] Posse S, Wiese S, Gembris D, Mathiak K, Kessler C, GrosseRuyken ML, Elghahwagi B, Richards T, Dager SR, Kiselev VG. Enhancement of BOLD-contrast sensitivity by single-shot multi-echo functional MR imaging. Magnetic resonance in medicine: official journal of the Society of Magnetic Resonance in Medicine / Society of Magnetic Resonance in Medicine 1999; 42:87–97.

[2] Poser BA, Versluis MJ, Hoogduin JM, Norris DG. BOLD contrast sensitivity enhancement and artifact reduction with multiecho EPI: parallel-acquired inhomogeneity-desensitized fMRI. Magnetic resonance in medicine: official journal of the Society of Magnetic Resonance in Medicine / Society of Magnetic Resonance in Medicine 2006; 55:1227–1235.

[3] Kettinger Á, Hill C, Vidnyánszky Z, Windischberger C, Nagy Z. Investigating the Group-Level Impact of Advanced Dual-Echo fMRI Combinations. Frontiers in neuroscience 2016; 10:571.

[4] Kundu P, Brenowitz ND, Voon V, Worbe Y, Vértes PE, Inati SJ, Saad ZS, Bandettini PA, Bullmore ET. Integrated strategy for improving functional connectivity mapping using multiecho fMRI. Proceedings of the National Academy of Sciences of the United States of America 2013; 110:16187–16192.

[5] Liu TT, Li B, Chen C, Fernandez B, Yang B, Banerjee S. Temporal SNR in multiecho fMRI and its dependence on the choice of weights. In: Proceedings of the 28th Annual Meeting of the ISMRM, Paris, 2020.

[6] Brennan LE, Reed LS. Theory of Adaptive Radar. IEEE Transactions on Aerospace and Electronic Systems 1973; AES-9:237–252.

[7] Monzingo R, Haupt R, Miller T, “Introduction to Adaptive Arrays”. Scitech Publishing, Raleigh, NC, 2nd ed., 2011.

[8] Jarrett D, Habets E, Naylor P, “Theory and Applications of Spherical Microphone Array Processing”. Springer, Switzerland, 2017.

[9] Gowland PA, Bowtell R. Theoretical optimization of multi-echo fMRI data acquisition. Physics in medicine and biology 2007; 52:1801–1813.

[10] Puckett AM, Bollmann S, Poser BA, Palmer J, Barth M, Cunnington R. Using multi-echo simultaneous multi-slice (SMS) EPI to improve functional MRI of the subcortical nuclei of the basal ganglia at ultra-high field (7T). NeuroImage 2018; 172:886–895.

[11] Cohen AD, Nencka AS, Lebel RM, Wang Y. Multiband multi-echo imaging of simultaneous oxygenation and flow timeseries for resting state connectivity. PLoS ONE 2017; 12:e0169253.

[12] Parrish TB, Gitelman DR, LaBar KS, Mesulam MM. Impact of signal-to-noise on functional MRI. Magnetic Resonance in Medicine 2000; 44:925–932.

[13] Krüger G, Glover GH. Physiological noise in oxygenation-sensitive magnetic resonance imaging. Magnetic Resonance in Medicine 2001; 46:631–637.

